# High-Frequency Spatial Feature Fusion with 3D CNN for Early Stage Schizophrenia Classification

**DOI:** 10.64898/2026.06.15.732490

**Authors:** Kamil Akhtar, Amirthavarshani Mahadevan

## Abstract

Early detection of schizophrenia (SZ) remains challenging due to the subtlety of early-stage brain alterations and reliance on subjective clinical assessment. We propose a frequency-aware 3D convolutional neural network (CNN) pipeline that integrates NeuroMark-HiFi high-pass spatial filtering with a modified VGGNet3D architecture featuring 3D Laplacian kernel initialization and dilated convolutions. Using the FBIRN dataset (N=311; 150 healthy controls, 161 SZ) with all 53 intrinsic connectivity networks (ICNs) per subject, we evaluate four experimental conditions across two hyperparameter configurations to isolate the contributions of enhanced input representations and frequency-aware model design. Under the optimized configuration, Condition 3 (HiFi + Laplacian initialization) achieved the best mean test accuracy of 75.54% with a peak single-fold accuracy of 87.10%, representing a 5.44% absolute gain over the optimized baseline. These results demonstrate that high-frequency spatial features are more discriminative for SZ classification than raw intensities, and that aligning Laplacian-initialized kernels with HiFi-filtered input creates a beneficial inductive bias—even with a compact model of approximately 1.4M parameters.

## I. Introduction

Deep learning, particularly Convolutional Neural Networks (CNNs), has markedly advanced medical image analysis by facilitating automated extraction of spatially meaningful representations from high-dimensional neuroimaging data. CNN adoption in medical image classification has risen from approximately 9% of published studies in 2014 to over 50% by 2023 [1], yet model performance remains sensitive to architectural design choices.

Functional magnetic resonance imaging (fMRI) has emerged as a key modality for studying brain functional connectivity. However, many approaches rely on original spatial representations, making detection of subtle brain network alterations, particularly in early-stage schizophrenia (SZ), challenging. SZ is a chronic brain disorder that disrupts perception, thinking, and behavior, with diagnosis often delayed by years due to lack of objective biomarkers. Conventional independent component analysis (ICA) methods lack sensitivity to fine-grained spatial variations within functional brain networks.

In this paper, we propose a frequency-aware 3D CNN pipeline combining high-pass filtered ICA representations with a modified VGGNet3D equipped with Laplacian kernel initialization and dilated convolutions. Our hypothesis is that integrating 3D high-pass filtering with a frequency-aware 3D-CNN will unlock discriminative high-frequency signals invisible to standard models. We evaluate this using the FBIRN dataset across four conditions and two hyperparameter configurations.

## II. Related Work

### A. Independent Component Analysis and Network-Level Insights

ICA has been widely used to extract spatially independent brain networks from resting-state fMRI (rs-fMRI) data. Traditional ICA methods, such as those in the GIFT toolbox [2], capture large-scale network activity but are limited in detecting high-spatial-frequency details reflecting fine-grained neurobiological abnormalities. Behzadfar et al. [3] introduced *NeuroMark-HiFi*, a frequency-informed ICA framework incorporating a 3D high-pass filter that improved detection of clinically relevant brain network alterations in SZ.

A key challenge with standard blind ICA is the correspondence problem: components from different subjects may not map to the same networks. Group Information Guided ICA (GIG-ICA) addresses this using a reference spatial template, ensuring cross-subject component correspondence [4]. We use the NeuroMark 1.0 template with 53 components spanning seven functional domains: Subcortical (SC), Auditory (AU), Sensorimotor (SM), Visual (VI), Cognitive Control (CC), Default Mode (DM), and Cerebellar (CB).

### B. Deep Learning in Functional Neuroimaging

3D CNNs have been effective for classifying neurological and psychiatric disorders from neuroimaging data, often outperforming classical machine learning. Qureshi et al. [5] demonstrated a modified 3D VGG-Net achieving 98.09% ±1.01% accuracy on the COBRE dataset for SZ classification, identifying visual-frontal disconnection and altered DMN-cerebellar connectivity. Dilated (atrous) convolutions have further improved CNN performance by expanding the receptive field without increasing parameters. Mahadevan et al. [6] showed that Laplacian-initialized kernels achieved 93.75% on fBIRN with an 18.75% improvement over random initialization, confirming dilation rate *d* = 3 as optimal. Their visualizations showed that Laplacian-initialized kernels produce sharper, more localized high-frequency activations.

### C. Bridging Frequency-Aware ICA and CNNs

Few studies have integrated frequency-informed ICA with CNN-based models for SZ classification. Most work focuses on either network-level connectivity or CNNs without spatial frequency consideration. Our study bridges this gap by combining NeuroMark-derived ICNs with 3D CNN architectures optimized for high-frequency feature extraction, unifying ICA interpretability with CNN predictive strength.

## I. Methods

To evaluate our hypothesis, we developed an experimental pipeline where the model takes ICNs as input and discriminates between SZ and healthy controls (HC).

### A. Data

We utilize data from the Functional Biomedical Informatics Research Network (FBIRN), a multi-site, multi-scanner fMRI dataset [7]. Phases II and III include individuals diagnosed with schizophrenia or schizoaffective disorder along with demographically matched healthy controls collected across multiple sites. The dataset consists of 150 healthy controls and 161 schizophrenia subjects (N=311 total), forming a relatively balanced cohort.

### B. Data Preprocessing

The fMRI data underwent an eight-step preprocessing pipeline: (1) variance normalization to standardize voxel intensities, (2) z-scoring to zero mean and unit variance, (3) motion regression using 6 rigid-body motion parameters, (4) 3rd-order polynomial detrending to remove slow signal drifts, (5) despiking to identify and correct outlier timepoints using Median Absolute Deviation (MAD *>* 2.5*σ*) with spline interpolation, (6) band-pass filtering to preserve temporal structures of interest, (7) final z-scoring to re-standardize after filtering, and (8) resampling to a uniform 2s sampling rate using rational resampling.

### C. Network Estimation

Following preprocessing, we applied the NeuroMark pipeline, a fully automated spatially constrained ICA framework [4], using the NeuroMark fMRI 1.0 template. This process yielded 53 intrinsic connectivity networks (ICNs) per subject as 3D spatial maps (dimensions: 53× 63× 52), along with corresponding 2D time courses. The 53 ICNs were grouped into seven functional domains: Subcortical (SC), Auditory (AU), Sensorimotor (SM), Visual (VI), Cognitive Control (CC), Default Mode (DM), and Cerebellar (CB). A sliding window approach with window size of 60 and step size of 1 was applied for temporal segmentation. For classification, all 53 component spatial maps were stacked as input channels to the 3D CNN, resulting in an input tensor of shape 53×63×52 per subject.

### D. High-Frequency Filtering (NeuroMark-HiFi)

To enhance sensitivity to fine-grained spatial features, we applied a 3D high-pass spatial filter to the ICN spatial maps following the NeuroMark-HiFi approach [3]. This filter operates in the frequency domain with a cutoff radius of 7, suppressing low-frequency structures while preserving high-spatial-frequency patterns. These high-frequency patterns represent subtle connectivity disruptions at network boundaries— precisely the kind of fine-grained alterations that are expected to differ between SZ and HC groups but are typically smoothed over by conventional analysis methods.

### E. Model Architecture

Our model is a modified VGGNet3D adapted from Qureshi et al. [5], with key modifications for frequency-aware processing and overfitting prevention. The architecture (Fig. 1) consists of three convolutional blocks followed by a Global Average Pooling (GAP) classification head.

**Fig. 1.**
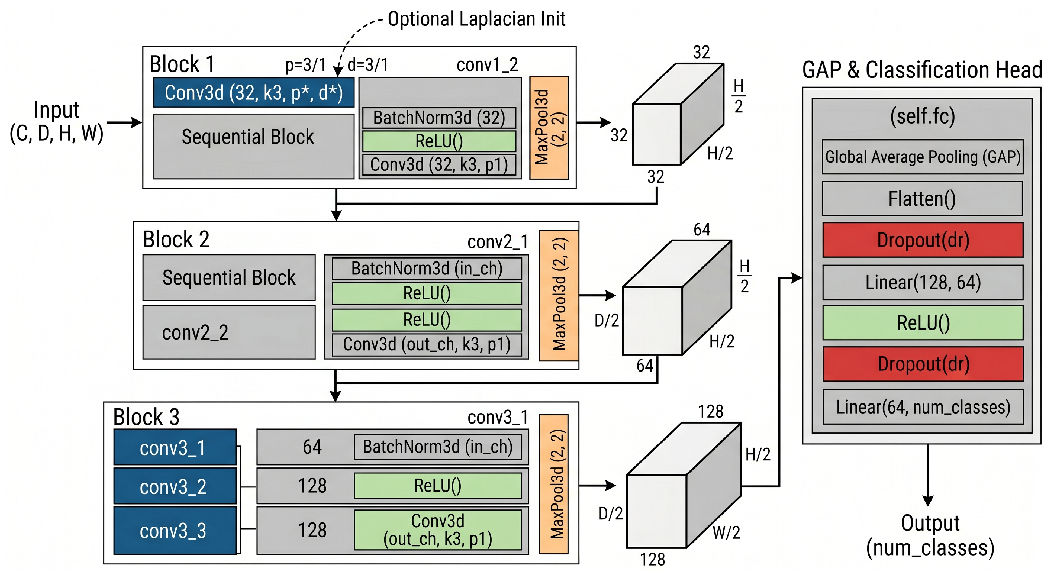
Modified VGGNet3D architecture overview. The model consists of three convolutional blocks with increasing filter depth (32→ 64→ 128), followed by a Global Average Pooling (GAP) classification head. The first convolutional layer optionally uses Laplacian initialization and dilated convolutions.

#### Block 1

The first layer is a Conv3D layer mapping the input channels (53 ICNs) to 32 filters with 3× 3× 3 kernels. When Laplacian initialization is enabled, the first 4 filters of this layer are pre-seeded with a normalized 3D Laplacian kernel. The discrete 3D Laplacian kernel assigns a center value of 26 with all 26 face, edge, and corner neighbors set to −1:

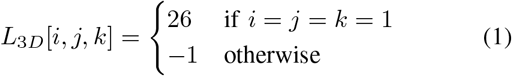

where indices are on a 3× 3× 3 grid. The kernel is L2-normalized before assignment, and small additive Gaussian noise (*σ* = 0.01) is applied for symmetry breaking. When dilation is enabled, this layer uses dilation rate *d* = 3 with padding=3, expanding the effective receptive field to 7× 7× 7 without increasing parameter count. The second layer in Block 1 applies BatchNorm → ReLU→ Conv3D (32→ 32, kernel 3× 3× 3), followed by 2× 2× 2 max-pooling.

#### Block 2

Two sequential BN → ReLU→ Conv3D layers transition from 32 to 64 filters, followed by max-pooling.

#### Block 3

Three sequential BN → ReLU → Conv3D layers operate at 128 filters (64→ 128, 128→ 128, 128 → 128), followed by max-pooling.

#### Classification Head

A critical modification from the original VGG architecture is the replacement of the traditional flatten-and-fully-connected head with: AdaptiveAvgPool3D(1) → Flatten→ Dropout → FC(128→ 64) → ReLU → Dropout→ FC(64→ 2). This GAP-based head reduces parameters by approximately 97.6% compared to a traditional dense head, eliminating the primary overfitting pathway while maintaining classification capacity. The total model size is approximately 1.2M parameters.

Training uses the Adam optimizer with cosine annealing learning rate scheduling and CrossEntropy loss. Evaluation is performed using 10-fold cross-validation with batch size of 12 and seed 28.

### F. Experimental Design

To systematically evaluate the contributions of input enhancement and architectural modifications, we designed four experimental conditions:

- **C1 (Baseline):** Raw ICA data + standard VGGNet3D (random init, no dilation).
- **C2 (HiFi Only):** HiFi-filtered data + standard VG-GNet3D.
- **C3 (HiFi + Laplacian):** HiFi-filtered data + Laplacian initialization, no dilation.
- **C4 (Full Stack):** HiFi-filtered data + Laplacian init + dilated convolutions (*d* = 3).

Each condition was run under two hyperparameter configurations. The *initial configuration* used: learning rate 0.001, weight decay 0.001, dropout 0.3, and 50 training epochs. The *optimized configuration* used: learning rate 0.0001, weight decay 0.005, dropout 0.5, and 100 training epochs. Both configurations used batch size 12 and 10-fold cross-validation.

## IV. Results

### A. Initial Configuration Results

Table I summarizes classification performance under the initial hyperparameter setting across all four conditions using all 53 ICNs per subject.

**TABLE 1.** Classification Results — Initial Configuration (53 ICNS)

| Condition | Mean Acc | Std Dev | Acc Range |
| --- | --- | --- | --- |
| C1: Baseline | 61.71% | 8.35% | 45.16–74.19% |
| C2: HiFi | 66.54% | 8.30% | 51.61–80.65% |
| C3: HiFi+Lap | **66.60%** | 9.58% | 51.61–87.10% |
| C4: Full Stack | 63.06% | 10.51% | 48.39–80.65% |

Under initial settings, the baseline already performed above chance at 61.71%, indicating that raw ICA spatial maps on 53 components contain some discriminative signal even without frequency enhancement. HiFi filtering improved mean accuracy by approximately 5% (C2: 66.54%), confirming that high-frequency spatial features carry additional discriminative information. Adding Laplacian initialization (C3: 66.60%) provided a marginal further gain in mean accuracy but notably achieved the highest single-fold accuracy of 87.10%, suggesting that on certain data splits, the Laplacian inductive bias captures strong discriminative patterns. The full stack condition (C4: 63.06%) unexpectedly underperformed C2 and C3, with the highest standard deviation (10.51%), indicating unstable training likely caused by the dilation expanding the receptive field too aggressively under the higher learning rate.

### B. Optimized Configuration Results

Table II presents results after hyperparameter tuning with lower learning rate, higher weight decay, stronger dropout, and extended training duration.

**TABLE 2.** Classification Results — Optimized Configuration (53 ICNS)

| Condition | Mean Acc | Std Dev | Max Acc |
| --- | --- | --- | --- |
| C1-Opt: Baseline | 70.10% | 5.93% | 80.65% |
| C2-Opt: HiFi | 73.64% | 6.90% | 83.87% |
| C3-Opt: HiFi+Lap | **75.54%** | 8.21% | **87.10%** |
| C4-Opt: Full Stack | 72.05% | 7.65% | 87.10% |

The optimized configuration substantially improved all conditions. The baseline jumped from 61.71% to 70.10%, confirming that the initial configuration was undertrained. Condition 3 (HiFi + Laplacian initialization) achieved the best mean accuracy of **75.54%** with a peak fold accuracy of **87.10%**, representing a 5.44% absolute gain over the optimized baseline and a 13.83% gain over the initial baseline. The optimized full stack (C4-Opt: 72.05%) improved considerably over its initial counterpart but still underperformed C3-Opt, confirming that dilation at *d* = 3 introduces excessive receptive field expansion when operating on 53 stacked ICN channels simultaneously.

### C. Key Observations

Across both configurations, several consistent patterns emerged. First, HiFi filtering consistently improved accuracy over raw baselines (+4.83% initial, +3.54% optimized), confirming that high-frequency spatial features carry discriminative information that is obscured in the raw data. Second, Laplacian initialization provided gains on top of HiFi filtering in both settings, validating the inductive bias created by aligning kernel initialization with input frequency content. Third, the full stack condition with dilation consistently underperformed Condition 3, indicating that the expanded 7× 7 ×7 receptive field may oversmooth features when all 53 components are used as input channels. Fourth, the optimized configuration reduced standard deviations across conditions (e.g., C1: 8.35%→ 5.93%), indicating more stable convergence with lower learning rate and stronger regularization.

## V. Discussion

The success of Condition 3 can be attributed to feature alignment: the Laplacian-initialized kernels are pre-tuned to detect high-frequency spatial boundaries, directly matching the frequency content of HiFi-filtered input data. This creates an inductive bias that accelerates learning and improves final performance without the instability introduced by dilated convolutions. Rather than learning to detect high-frequency patterns from scratch, the network begins with kernels already tuned to the relevant frequency band.

The underperformance of Condition 4 (full stack with dilation) across both configurations was unexpected given prior results on individual components [6]. However, this can be explained by the difference in input dimensionality. When operating on 53 stacked ICN channels, the network already receives rich spatial context from multiple brain networks simultaneously. The dilated convolutions with *d* = 3 expand the receptive field to 7 ×7× 7, which may cause oversmoothing of the fine-grained boundary features that the HiFi filter was designed to preserve. In contrast, when processing individual components (as in [6]), the dilation provides necessary spatial context that is otherwise absent.

The GAP head replacement was critical for training on N=311 subjects. By reducing parameters by 97.6% compared to traditional fully-connected layers, the model avoids the overfitting that commonly affects large architectures on small neuroimaging datasets. The improvements from the optimized configuration (learning rate: 0.001→ 0.0001, dropout: 0.3→ 0.5, weight decay: 0.001→ 0.005, epochs: 50→ 100) further highlight the sensitivity of small-sample deep learning to regularization strategy and training duration.

Compared to Qureshi et al. [5] who achieved 98.09% on the COBRE dataset (N=144), our lower absolute accuracy reflects the multi-site nature of FBIRN introducing scanner variability, and the use of all 53 ICNs rather than selected discriminative components. However, our best single-fold accuracy of 87.10% suggests that with better component selection and stronger regularization, substantially higher mean accuracy is achievable.

Several limitations should be noted. The overall accuracy of 75.54% is still below clinical utility thresholds. Using all 53 ICN components may dilute discriminative signal with non-informative networks. Additionally, validation on datasets beyond FBIRN would be needed to assess generalizability.

## VI. Conclusion

We demonstrated that high-frequency spatial features from NeuroMark-HiFi filtering are more discriminative for SZ classification than raw ICA intensities. HiFi + Laplacian initialization (C3) achieved the best performance at 75.54% mean accuracy (87.10% peak), validating a pipeline bridging frequency-aware signal processing with domain-informed deep learning. The key finding is that frequency alignment between input and model initialization creates a beneficial inductive bias, while excessive receptive field expansion through dilation can be counterproductive when multiple ICN channels already provide sufficient spatial context. Future work will focus on selecting a subset of discriminative components and exploring alternative dilation strategies.

